# Comprehensive transcriptional profiling and mouse phenotyping reveals dispensable role for adipose tissue selective long noncoding RNA *Gm15551*

**DOI:** 10.1101/2021.12.17.473160

**Authors:** Christoph Andreas Engelhard, Chien Huang, Sajjad Khani, Petr Kasparek, Jan Prochazka, Jan Rozman, David Pajuelo Reguera, Radislav Sedlacek, Jan-Wilhelm Kornfeld

## Abstract

Cold and nutrient activated brown adipose tissue (BAT) is capable of increasing systemic energy expenditure via uncoupled respiration and secretion of endocrine factors thereby protecting mice against diet-induced obesity and improving insulin response and glucose tolerance in men. Long non-coding RNAs (lncRNAs) have recently been identified as fine tuning regulators of cellular function. While certain lncRNAs have been functionally characterised in adipose tissue, their overall contribution in the activation of BAT remains elusive. We identified lncRNAs correlating to inter-scapular brown adipose tissue (iBAT) function in high fat diet (HFD) and cold stressed mice. We focused on *Gm15551* which has an adipose tissue specific expression profile, is highly upregulated during adipogenesis and downregulated by β-adrenergic activation in mature adipocytes. Albeit we performed comprehensive transcriptional and adipocyte physiology profiling *in vitro* and *in vivo*, we could not detect an effect of gain or loss of function of *Gm15551*.

## 4 Introduction

The prevalence of obesity is increasing worldwide (NCD Risk Factor Collaboration (NCD-RisC), 2016). Obesity is the result of a chronic imbalance between energy intake and expenditure resulting in the accumulation of excess adipose tissue. Obesity is correlated with increased overall mortality and is a risk factor for various diseases including cardio vascular disease and diabetes type 2 (Angelantonio et al., 2016; Prospective Studies Collaboration, 2009).

Adipose tissue plays a central role in the regulation of energy balance. While white adipose tissue (WAT) mainly functions as storage of excess energy in the form of triglycerides, BAT is a highly metabolically active tissue (Rosen and Spiegelman, 2014). Morphologically, BAT is densely packed with mitochondria and generates heat by short-circuiting the mitochondrial proton gradient via uncoupling protein 1 (*UCP1*), facilitating substrate use without ATP generation (Cannon and Nedergaard, 2004). Thereby active BAT significantly improves glucose and lipid clearance and raises energy expenditure (Betz and Enerbäck, 2017; Klepac et al., 2019). Additionally, active BAT signals to other tissues improving the whole body metabolic profile via the secretion of endocrine factors and micro RNA (miRNA) containing exosomes (Scheele and Wolfrum, 2020; Zhang et al., 2019). In this regard, the recent demonstration of the presence of active BAT in adult humans has led to an increased interest in understanding the molecular signals underlying BAT differentiation and function (Betz and Enerbäck, 2017; Nedergaard et al., 2007).

The characterisation of the human transcriptome in the course of the ENCODE project revealed pervasive transcription of three quarters of the human genome (Djebali et al., 2012). Most of the transcribed sequences however do not fall within protein coding regions but give rise to non-coding ribonucleic acid (RNA) such as lncRNA (Djebali et al., 2012). lncRNAs are defined as non-coding genes giving rise to transcripts of more than 200 nt, which do not belong to an otherwise functionally defined class of RNA (Gil and Ulitsky, 2019). The lack of a functional definition coincides with a broad range of modes of function: lncRNAs have been shown to act both *in cis* as well as *in trans* (Gil and Ulitsky, 2019; Yao et al., 2019) via an interaction of the transcribed RNA molecule with other RNA, proteins or the DNA (Nguyen et al., 2018; Yi et al., 2020). Compared to coding genes, lncRNAs are on average lower expressed but show more tissue and developmental stage specific expression profiles, advocating for a role as fine tuning regulators of cellular function (Derrien et al., 2012). Selected lncRNAs have been shown to interfere with adipose tissue function and differentiation such as *lncBATE10* which acts as a decoy for Celf1 which would otherwise bind to and repress Pgc1a mRNA (Bai et al., 2017), *H19* which functions as a BAT-specific gatekeeper of paternally expressed genes (Schmidt et al., 2018) and *Ctcflos* which regulated expression and splicing of *Prdm16* (Bast-Habersbrunner et al., 2021). However, their overall contribution to these processes remains elusive.

In this study, we performed RNA sequencing on BAT from C57BL/6 mice challenged with cold-treatment and high-fat diet, two physiologically relevant models of BAT activation (Alcalá et al., 2017; Cannon and Nedergaard, 2004) as well as on a set of seven metabolically active tissues. We found a set of adipose tissue specific cold and/or diet regulated lncRNAs, from which we selected *Gm15551* as a candidate for functional studies. The genomic locus of *Gm15551* is bound by Pparg and Prdm16 in brown adipocytes and it is upregulated in adipogenesis, and downregulated upon β-adrenergic stimulation in adipocytes. We performed comprehensive phenotyping of gain and loss of function *in vitro* as well as loss of function *in vivo* but could not detect any phenotype related to *Gm15551*.

## 5 Results

### Total RNA-seq identifies lncRNAs regulated in activated iBAT

In order to identify lncRNAs implicated in the regulation of iBAT function we set out to perform total RNA-seq on C57BL/6N mice put on a high fat diet regime from 8 weeks of age onwards for 12 weeks and additionally housed at 4 °C for 24 h at the end of this period (Fig 1A). We found that cold treatment significantly reduced the weight of the iBAT in both the control and the HFD group (Fig S1A), while HFD treatment alone did not induce significant changes in iBAT weight. On the other hand, epididymal white adipose tissue (eWAT) and inguinal white adipose tissue (iWAT) as well as liver weights were increased upon HFD treatment, independent of the cold treatment. Cold treatment alone only induced an increase of eWAT but not iWAT and liver weight. Gene expression measurements reflected the observed difference in the reaction of white and brown adipose tissue to the treatments (Fig S1B). Cold treatment induced a robust induction of the common adipose marker gene *Elovl3* as well as the brown adipose markers *Cidea*, *Dio2* and *Ucp1* while HFD alone was insufficient for the induction of any significant changes in those genes, although *Ucp1* was tendentiously upregulated. In iWAT however, cold and HFD treatment showed opposing effects. The white adipose marker gene *Lep* was tendentiously repressed upon cold treatment and induced by HFD while *Elovl3*, *Dio2* and *Ucp1* were upregulated by the cold and downregulated by the HFD treatment. The treatment regimes also directly affected the animals’ metabolism as seen by the significantly impaired glucose tolerance upon HFD treatment (Fig S1C). Energy expenditure was elevated by cold treatment but reduced by HFD treatment (Fig S1D) while the respiratory exchange rates indicated a shift towards lipid catabolism induced by both cold and HFD treatment (data not shown). Together, these data indicate our dataset is an adequate model for different functional states of iBAT. Additionally, we used a second dataset consisting of seven metabolically active tissues (iBAT, iWAT, eWAT, liver, kidney, muscle, heart) which we have generated for a previous study to be able to assess transcriptome wide tissue specificity (Pradas-Juni et al., 2020). To generate a comprehensive set of lncRNA genes, we combined the annotated transcript isoforms from GENCODE and the lncRNA isoforms from RNAcentral on gene level.

**Fig 1:**
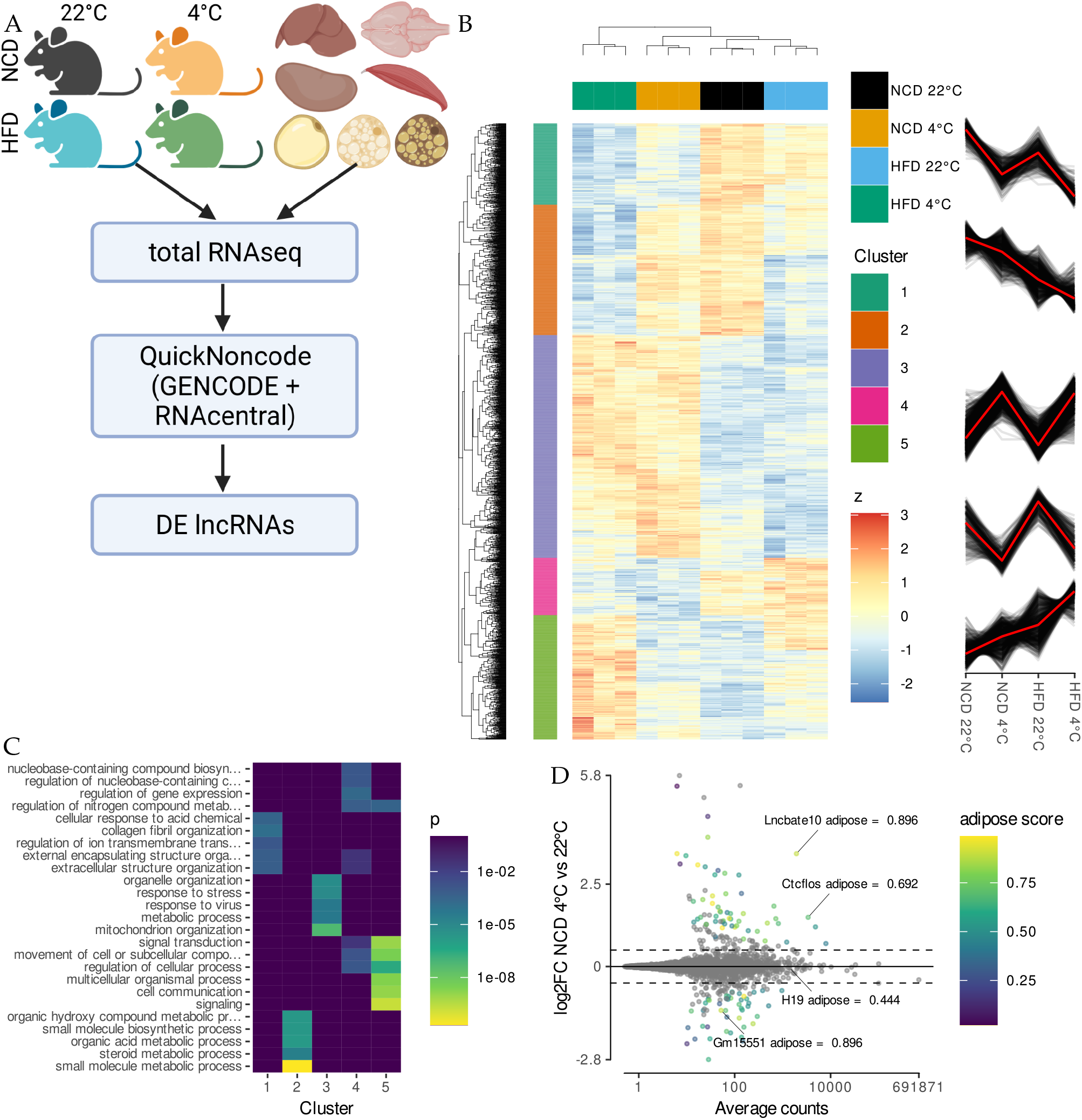
RNA-Seq reveals temperature and obesity dependent changes in iBAT lncRNA expression. **A** Experimental design. Total transcriptomes from the iBAT of 20 week old mice housed at 22 °C or 4 °C for 24 h (n = 3) fed either a high fat or a control diet for 12 weeks were analysed together with the total-transcriptomes from seven metabolically active tissues (n = 1, GSE121345). The union of GENCODE and RNAcentral annotated genes was used for the analysis to reveal lncRNAs which are both adipose tissue specific and regulated by physiologically relevant stimuli. Created with BioRender.com. **B** Hierarchical clustering of genes differentially regulated by diet and cold treatment in adipose tissue (likelihood ratio test, FDR < 0.001). Colour code depicts row wise standardised expression. **C** GO enrichment analysis for the gene clusters shown in B. **D** Expression levels and changes for lncRNA genes in iBAT from cold treated compared to control mice on control diet. Genes showing significant differential gene expression are colour coded indicating their adipose tissue specificity (wald test, log2 fold change (log2FC) > 0.5, n = 6, *s* < 0.05).

Total RNA-seq of the iBAT dataset identified 2490 differentially expressed genes, among them 216 lncRNA genes (likelihood ratio test (LRT), *p* < 0.001; Fig 1B). The largest cluster (cluster 3) consisted of genes that were induced by cold treatment independent of the diet and was enriched for genes involved in stress response and mitochondrion organisation (Fig 1C, Fig S1E). Similarly, cluster 4 contained genes downregulated by cold treatment independently of diet and was enriched for gene expression regulation and signal transduction. The other clusters included genes with synergistic interaction of HFD and housing temperature; either being induced (cluster 5, enriched for signalling) or repressed by HFD and cold treatment (clusters 1 and 2, enriched for extracellular matrix as well as metabolism). We used the transcriptomics data from the seven metabolically active tissues to calculate an adipose tissue enrichment score, defined as the ratio of log adipose tissue counts over the log of total counts. As temperature influenced the gene expression of more genes then HFD, we focused on genes regulated by the cold treatment. Wald tests identified 110 cold regulated lncRNA genes (*s* < 0.05), of which 65 (59 %) also showed an adipose tissue specific expression profile (adipose score > 50 %; Fig 1D, Fig S1G). On the other hand, from the 610 cold regulated coding genes, only 35 % showed adipose tissue specific expression (Fig S1F). Noteworthy, our analysis identified known brown adipose marker genes such as *Ucp1* and *Adcy3* as well as the lncRNA genes *LncBate10* and *Ctcflos*, which have previously been shown to play a role in regulation of brown/beige adipose tissue function (Bai et al., 2017; Bast-Habersbrunner et al., 2021), proving the applicability of our strategy towards the identification of novel candidate adipose regulating lncRNAs. In order to rule out that any of the identified lncRNA genes were differentially regulated because of an increased immune cell infiltration of the iBAT caused by the cold or HFD treatment (Alcalá et al., 2017), we checked the expression profiles of several immune cell marker genes (Henriques et al., 2020), of which none were differentially regulated (Fig S1H).

### *Gm15551* is an adipose specific, highly regulated lncRNA

We further focused our study on the lncRNA *Gm15551*, which was highly adipose specific and significantly repressed upon cold treatment in iBAT (Fig 1D). While HFD alone was not sufficient to induce the repression of *Gm15551* expression, the combination of cold treatment and HFD further repressed *Gm15551* compared to cold treatment alone (ANOVA; *p* = 0.000701, Fig 2A). Among the examined tissues, the expression of *Gm15551* was observed to be strictly restricted to adipose tissue, similar to the common adipocyte marker genes *Adipoq* and *Pparg* (Fig 2B). Within the three adipose tissues we looked at, *Gm15551* showed the highest expression in eWAT and the lowest in iBAT, with intermediate expression in iWAT, anti-correlating to the expression of the thermogenic adipocyte marker gene *Cidea*. The notion of this expression pattern together with the repression upon activation of thermogenesis in iBAT led us to hypothesize *Gm1555* might have an anti-thermogenic function.

**Fig 2:**
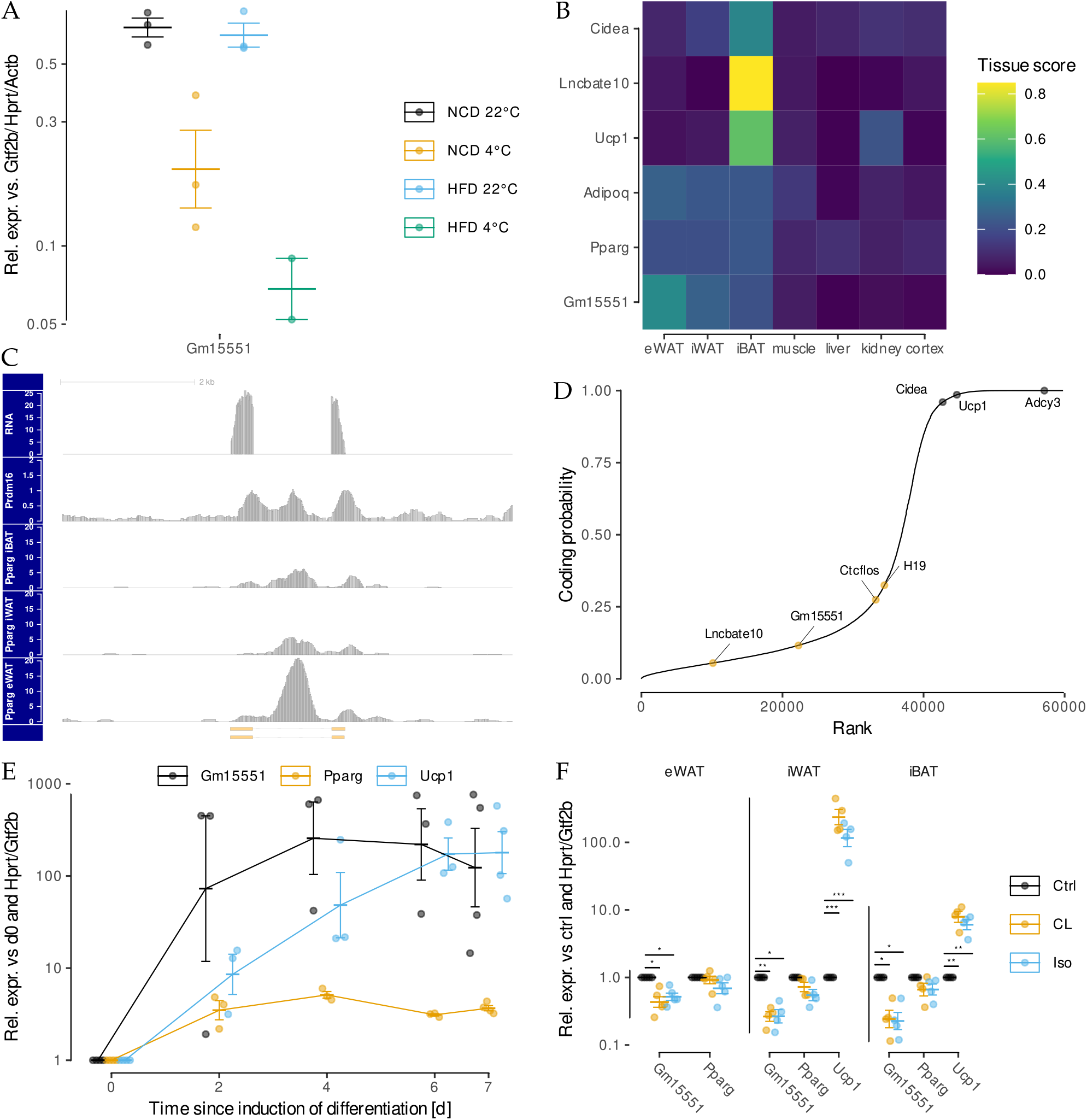
*Gm15551* is an adipose tissue specific, diet and temperature regulated lncRNA. **A** Expression of *Gm15551* in iBAT from cold treated and/or HFD fed mice. **B** Expression profile of *Gm15551*, the common adipocyte marker genes *Pparg* and *Adipoq* as well as the iBAT specific lncRNA *Lncbate10* and the brown adipocyte marker genes *Ucp1* and *Cidea* in seven metabolically active tissues. **C** Genomic locus of *Gm15551* showing the RNA expression as well as binding sites of *Prdm16* in iBAT (PRJNA269620) and *Pparg* in eWAT, iWAT and iBAT (PRJNA177164). **D** Ranked coding probability of all genes expressed in the dataset as calculated by CPAT. Indicated are *Gm15551*, the coding genes *Ucp1*, *Adcy3* and *Cidea* as well as the lncRNAs *Ctcflos*, *H19* and *LncBate10*. **E, F** Expression profiles of *Gm15551*, the common adipocyte marker gene *Pparg* and the brown adipocyte marker gene *Ucp1* during the differentiation of PIBA cells (E) and in fully differentiated primary adipocytes (F) stimulated for 24 h with the non-selective β-adrenergic agonist isoproterenol or the β_3_-specific agonist CL316243 (paired *t*-test, n = 4-5).

*Gm15551* is expressed from chromosome 3 and there are 2 transcripts annotated which both consist of 2 exons and only differ in the exact position of the transcription end site (Fig 2C). Analysis of publicly available ChIP-Seq data showed that the locus is bound by the core thermogenic transcription factor Prdm16 in iBAT. Additionally, we found that Pparg binds to the *Gm15551* locus in eWAT, iWAT as well as iBAT with the height of the ChIP-Seq peak correlating with the *Gm15551* RNA expression levels. Chromatin features such as the ratio of H3K4me1 relative to H3K4me3 have previously been used to distinguish promoters from enhancers (Natoli and Andrau, 2012). Therefore we looked at histone modifications using chromatin immunoprecipitation (ChIP)-Seq (S2A). We found higher levels of H3K4me3 compared to H3K4me1 which is indicative of a promoter as opposed to an enhancer. H3K327ac signal was higher then H3K4me3 or H3K4me1 and H3K327me3 was basically absent.

As *Gm15551* is expressed antisense from a locus within intron 2 of the intracellular Ca^2+^ signalling protein *Camk2d* and it is known that lncRNA can work as *in cis* regulators of nearby coding genes (Gil and Ulitsky, 2019), we checked whether the expression of *Gm15551* correlates with the expression of *Camk2d* in various publicly available RNA-Seq datasets of adipose tissue, but found no significant correlation (ANCOVA, *p* = 0.848; Fig S2B). To exclude the possibility of *Gm15551* being a coding gene wrongly annotated as lncRNA (Anderson et al., 2015), we calculated the coding potential for all genes expressed in our dataset using CPAT (Wang et al., 2013). *Gm15551* showed a low coding probability comparable to other known lncRNA genes as opposed to the brown adipocyte marker genes *Cidea*, *Adcy3* and *Ucp1* (Fig 2D). Similarly, ranking genes by the ratio of ribosome associated over total RNA in a publicly available TRAP-Seq data set of iBAT sorted *Gm15551* with other lncRNA genes (Fig S2C).

Next we followed the gene expression of *Gm15551* during the differentiation of preadipocytes into mature adipocytes. Our analysis showed that *Gm15551* is highly upregulated already early in differentiation, similar to the core adipocyte transcription factor *Pparg* and unlike *Ucp1* which only reaches maximum levels in late differentiation (Fig 2E). In order to mimic the effects of cold treatment on adipose tissue *in vitro*, we stimulated cells with the non-selective β-adrenergic agonist isoproterenol or β__3__ specific agonist CL316243. Both stimuli were sufficient to repress *Gm15551* in differentiated adipocytes originating from eWAT, iWAT and iBAT (Fig 2F). In primary immortalized brown adipocytes, the effect of β-adrenergic stimulation on the expression of *Gm15551* was stable over 24 h (Fig S2D).

### Gain- and loss-of-function of *Gm15551* does not disturb brown adipocyte development and function *in vitro*

To investigate the role of *Gm15551* in differentiation and function of brown adipocytes, we used an immortalized brown preadipocyte cell line stably expressing the CRISPRa SAM system for gain-of-function studies together with locked nucleic acid (LNA) antisense oligonucleotides for loss-of-function studies (Lundh et al., 2017). Transfection of either one of two plasmids encoding single guide RNAs (sgRNAs) targeting *Gm15551* two days prior to the induction of the differentiation led to a robust overexpression of *Gm15551* compared to the empty vector control at day 1 of differentiation (Fig 3A). The effect of the overexpression was greatly diminished on day 4 and 7 because the natural gene expression of *Gm15551* rises during differentiation. However, we could not observe any changes in the expression of common and brown adipocyte marker genes or in the cells ability to accumulate lipids (Fig 3A, B, Fig S3A).

**Fig 3:**
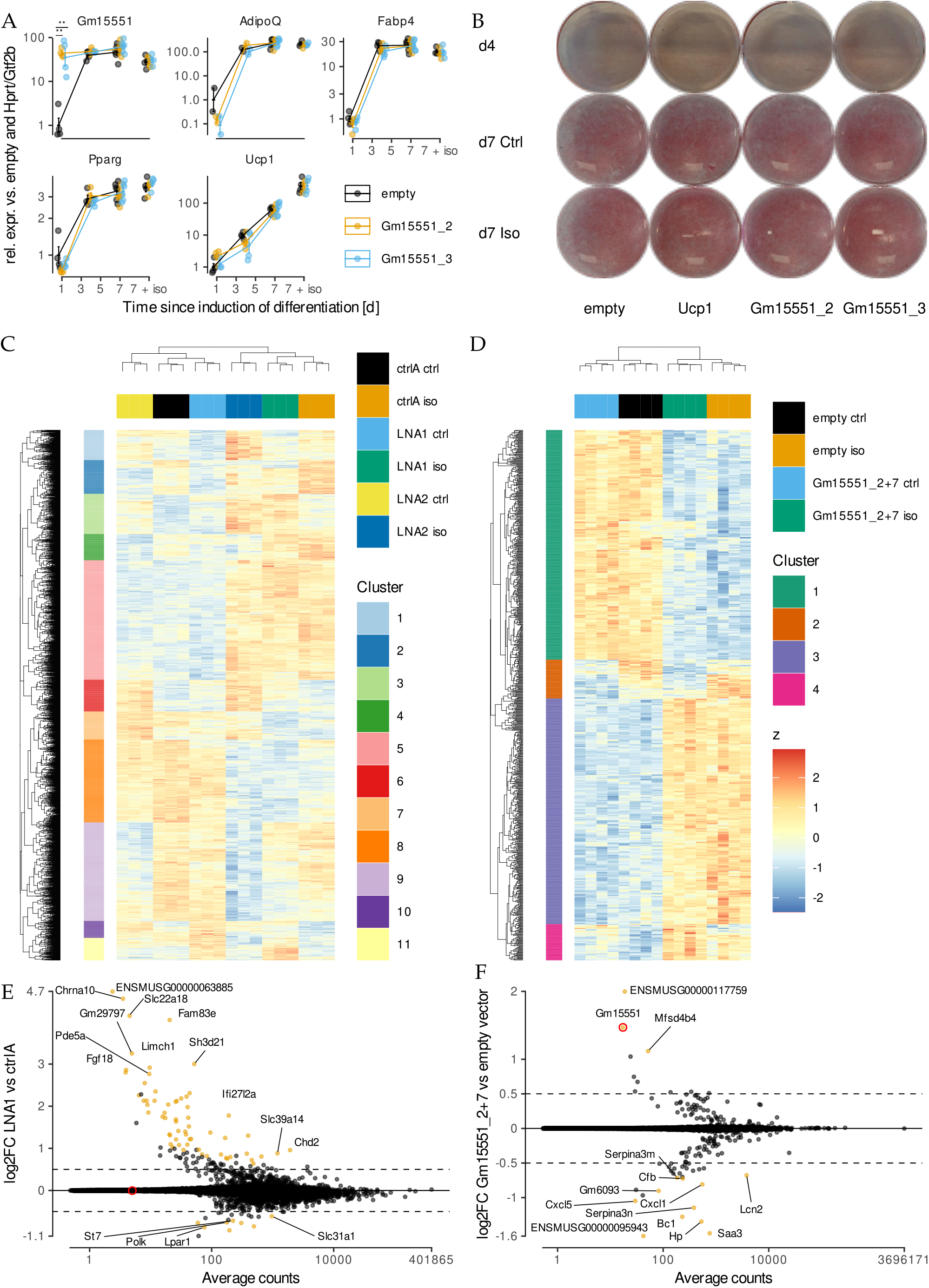
*Gm15551* is dispensable for iBAT function *in vitro*. **A, B** Gene expression profiles (A) of *Gm15551*, the common adipocyte marker genes *Pparg*, *Adipoq* and *Fabp4* as well as the brown adipocyte marker gene *Ucp1* (paired *t*-test, n = 2 - 6) and oil red o staining (F) of wt1-SAM cells transfected with plasmids coding for sgRNAs targeting *Gm15551*, *Ucp1* or empty vector two days before induction of differentiation. **C, D** Hierarchical clustering of genes differentially regulated by β-adrenergic stimulation and/or knockdown (C) or overexpression (D) in mature adipocytes at day 4 of differentiation (LRT, n = 3 (C) or 4 (D), *p* < 0.001). **E, F** Effect of knockdown (E) or overexpression (F) of *Gm15551* on gene expression in mature wt1-SAM cells. The log2FC is the average over the effect in isoproterenol stimulated and control cells (wald test, log2FC > 0.5, n = 6 (E) or 8 (F), *s* < 0.05).

Next we set out to knock down *Gm15551* in mature adipocytes. In order to detect potential interactions of *Gm15551* expression with thermogenic activation of brown adipocytes, we looked at cells both under basal conditions and under β-adrenergic stimulation. Reverse transfection with two different LNAs targeting *Gm15551* on day 4 of differentiation resulted in robust downregulation of *Gm15551* on day 7 compared to the non-targeting control LNA (Fig S3B). Overall, we found 2762 genes differentially regulated by either knockdown or stimulation (Fig 3C). Hierarchical clustering showed, that the influence of the β-adrenergic stimulation was more pronounced then that of the loss-of-function of *Gm155551*. Samples treated with the control non-targeting LNA clustered together with LNA1 treated samples in both the basal and stimulated condition, indicating that the two LNAs used caused different effects. Looking at the specific effect of the each LNA individually, we found 70 and 188 differentially regulated genes respectively (Fig 3E, Fig S3C). There was only an overlap of 16 genes detected to be significantly regulated by the knockdown of *Gm15551* using either of the two LNAs. Gene ontology (GO) analysis revealed that these genes were enriched for signalling and especially NF-kB mediated signalling (data not shown).

Similarly, reverse transfection with sgRNA encoding plasmids led to a small but significant over-expression of *Gm15551* in fully matured adipocytes and noteworthy was able to suppress its down-regulation upon β-adrenergic stimulation (Fig S3D). However, we did not observe any changes in the gene expression of any of the probed adipocyte marker genes. We further raised the overexpression efficiency by simultaneous transfection of two different plasmids encoding sgRNAs targeting *Gm15551* (Fig S3E). We sequenced the transcriptomes of these samples and found a total of 792 genes differentially regulated by either gain-of-function of *Gm15551* or β-adrenergic stimulation (Fig 3D). The effect of the thermogenic activation dominated the dataset as shown by hierarchical clustering. However, there were also two clusters with genes affected by the overexpression of *Gm15551*. When we specifically looked for changes in gene expression caused by the *Gm155551* gain-of-function, we found 14 differentially expressed genes and GO analysis showed an enrichment for genes involved in inflammatory response (Fig 3E, F).

Comparison between the genes differentially regulated by the gain and loss-of-function of *Gm15552* showed that there was no overlap. Additionally, most genes affected by the knock down of *Gm15551* showed no change in gene expression in the gain-of-function experiment (Fig S3H). Finally, Oil red O staining of mature adipocytes showed no effect of either gain or loss-of-function of *Gm15551* on the cells’ ability for lipid accumulation (Fig S3F).

### *Gm15551* loss-of-function does not impair adipose tissue function *in vivo*

Next we created a loss-of-function mouse model by knocking out exon 1 of *Gm15551*. Since brown adipose tissue plays a role in the regulation of body weight as well as lipid and glucose metabolism (Rui, 2017), we challenged homozygous ∆Gm15551 mice and wild type litter mates from 8 weeks of age for 12 weeks with a high fat diet and repeatedly measured body weight and performed intraperitoneal glucose tolerance test (IPGTT), and indirect calorimetry (Fig S4A). While the HFD was sufficient to provoke a significant raise in body weight (Fig 4A; *p* = 0.038) characterised by an increased amount of body fat (Fig S4B; *p* = 0.027), we did not observe significant changes induced by the loss-of-function of *Gm15551* (*p* = 0.29 and 0.76 respectively). Similarly, prolonged HFD treatment but not *Gm15551* loss-of-function resulted in impaired glucose tolerance (Fig 4B; no p as too little n). Additionally, adipocyte diameter and morphology of HFD treated animals was not affected by *Gm15551* knockout (Fig S4C, D; *p* = 0.359). Next we performed indirect calorimetry while sequentially changing the temperature first from room temperature to thermoneutrality (30 *◦*C), followed by a period at 4 *◦*C before returning to room temperature (23 *◦*C). Upon the beginning of thermoneutrality, energy expenditure slightly dropped and consequently raised when the temperature was dropped. Upon return to room temperature, the energy expenditure went back to the starting point (Fig 4C). However, there was no effect of the *Gm15551* knock out (no statistics done so far because to little number of animals). Respiratory exchange ratio of control diet animals raised with the onset of the first dark phase at thermoneutrality and dropped again in the following light phase indicating combustion of carbohydrates taken up with the food during dark phase(Fig 4D). With prolonged cold treatment, the respiratory exchange rate rose again to an intermediate value indicating the mice had to take up food in addition to combusting stored lipids. At the end of the cold treatment, the respiratory exchange rate rose even further indicative of the mice mostly relying on energy from the taken up carbohydrates. This effect of temperature and day light cycle was mostly suppressed in HFD animals as they take up less carbohydrates with their alimentation. Again there was no evidence of an impact of the *Gm15551* loss-of-function (no statistics done).

**Fig 4:**
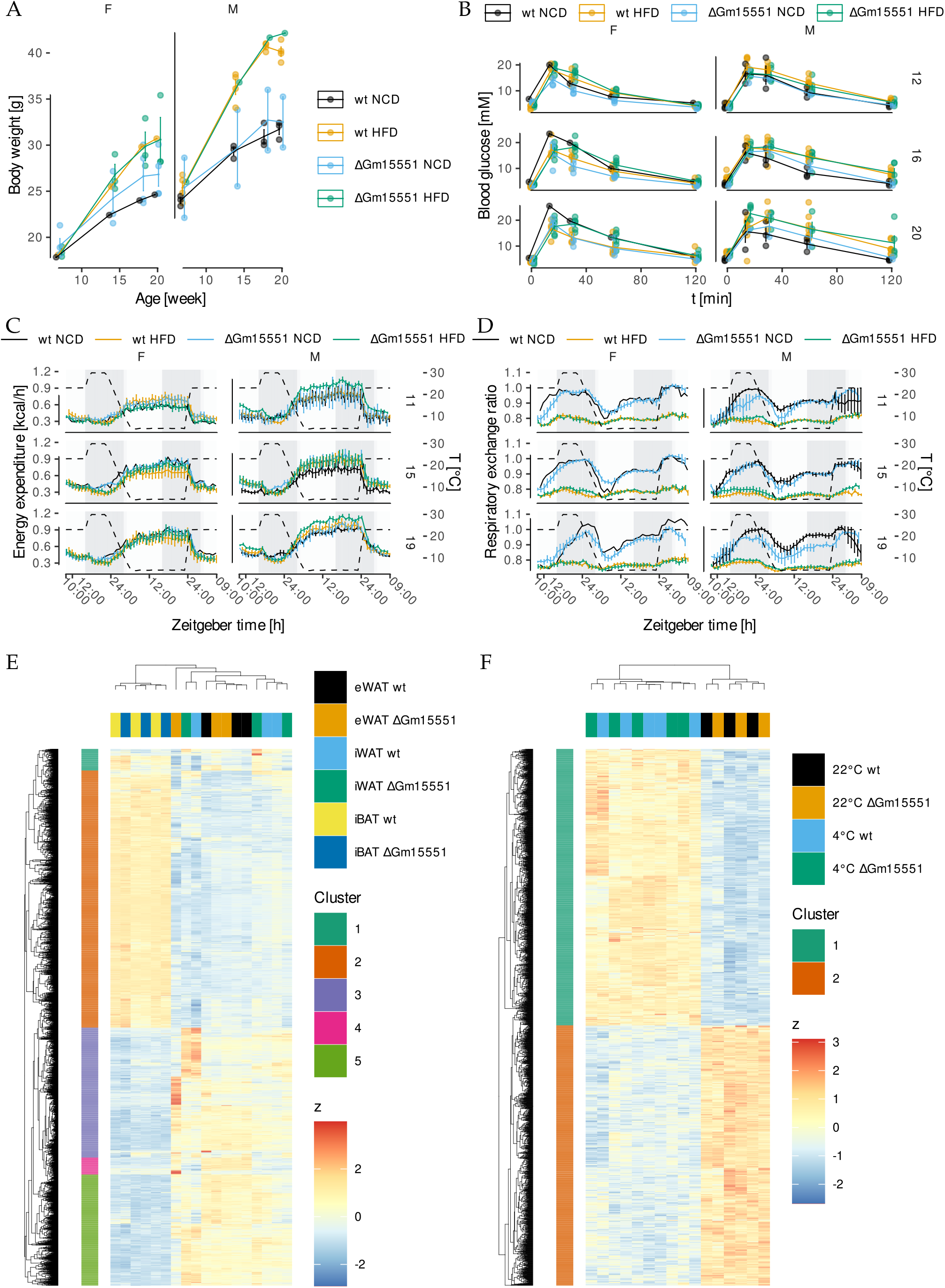
*Gm15551* is dispensable for iBAT function *in vivo*. **A** Body weight of ΔGm15551 and wild type mice fed a high fat or control diet. **B** IPGTT of 12, 16 and 20 week old ΔGm15551 and wild type mice fed a high fat or control diet. **C, D** Energy expenditure (C) and respiratory exchange rates (D) of 12, 16 and 20 week old ΔGm15551 and wild type mice fed a high fat or control diet. **E** Hierarchical clustering of genes differentially regulated between adipose tissues or by knockout of *Gm15551* in 12 week old mice (LRT, n = 3, *p* < 0.001). **F** Hierarchical clustering of genes differentially regulated by temperature or by knockout of *Gm15551* in in iBAT of 12 week old mice (LRT, n = 3 or 5, *p* < 0.001).

When we compared the adipose tissue transcriptomes from HFD and control diet fed animals, we found 5655 differentially expressed genes (LRT, *p* < 0.001; Fig 4C). Hierarchical clustering was strongly driven by the difference between brown and white adipose tissues. Gene ontology analysis showed that the genes with higher expression in brown adipose tissue were enriched for the terms related to mitochondria, while genes showing a higher expression in white adipose tissue were enriched for terms related to immune system and locomotion (Fig S4E). The direct comparison of the samples from wild type animals with those from knock out animals revealed only 11 differentially regulated genes (wald test, *s* < 0.05; S4F). Additionally, we compared the gene expression for iBAT from wild type and ∆Gm15551 mice both at room temperature and after 24 h of cold treatment. Overall, there were 2531 differentially expressed genes (LRT, *p* < 0.001), which clustered the samples by temperature but not genotype (4F). Both sets of up and down regulated genes upon cold treatment showed enrichment for terms related to metabolism (S4G). Also the direct comparison of the wild type transcriptomes with those from ∆Gm15551 animals only revealed 6 differentially expressed genes (wald test, *s* < 0.05; S4H).

## 6 Discussion

Cold and nutrient activated BAT regulates energy homeostasis and improves metabolic status via non-shivering thermogenesis and the secretion of endocrine factors (Betz and Enerbäck, 2017; Scheele and Wolfrum, 2020). lncRNAs have been shown to be tissue specific fine tuning regulators of tissue function, and therefore have been proposed as potential selective targets for the treatment of different diseases (Matsui and Corey, 2017; Wahlestedt, 2013). In the recent years, the function of some lncRNAs expressed in adipose tissue has been described (reviewed by Sun and Lin, 2019). However, the function of most lncRNAs remains unknown. Here, we detected a set of 65 lncRNAs whose expression is specific to adipose tissue and correlates with BAT function and characterised *Gm15551* further *in vitro* and *in vivo*.

We found *Gm15551* to be highly adipose tissue specific with a higher expression in white compared to brown adipose tissues. *Gm15551* is highly induced in the early stages of brown adipogenesis and downregulated upon beta adrenergic stimulation in both white and brown adipocytes. We could show that the key transcription factor Prdm16, which controls the determination of brown adipocytes and the browning of white adipose tissue (Seale et al., 2011, 2007), binds to the *Gm15551* locus in iBAT. Further, we found the *Gm15551* locus to be occupied by Pparg in white as well as brown adipose tissues with a more pronounced occupancy in white compared to brown adipose tissue. Pparg is a transcription factor involved in the maintenance of the general adipocyte phenotype but also showing depot specific binding patterns (Siersbæk et al., 2012). These findings led us to hypothesize an adipocyte specific function of *Gm15551*.

Previous studies have shown that lncRNAs might give rise to unidentified translation products (Anderson et al., 2015; Ji et al., 2015). We used a sequence based bioinformatics tool to calculate coding probability and analysed a public TRAP-Seq data set to detect ribosome associated RNAs. Our results showed that *Gm15551* has a low coding probability and is not associated with ribosomes. While enhancers are known to give rise to bidirectionally transcribed, short, unspliced and unstable enhancer RNAs (eRNAs), it has recently been reported that some enhancers can also be the place of unidirectional transcription giving rise to spliced lncRNAs (Gil and Ulitsky, 2018; Natoli and Andrau, 2012). The *Gm15551* locus featured a low ratio of the H3K4me1 over the H3K4me3 histone mark, indicative of promoters. However, the *Gm15551* locus also features H3K27ac histone marks, high levels of which are characteristic for enhancers (Natoli and Andrau, 2012). Enhancers are cis-regulatory elements, regulating the expression of nearby genes. Therefore we compared the expression of *Gm15551* and *Camk2d*, which overlaps with *Gm15551* in the genome, in several adipocyte related RNA-Seq datasets but found no correlation. However, we cannot rule out a potential enhancer function of *Gm15551*.

In order to unveil potential effects of *Gm15551* gain-of-function on brown adipogenesis, we over-expressed *Gm15551* two days prior to the induction of differentiation in a brown preadipocyte cell line, but could not measure any effect of the overexpression neither on common and brown adipocyte marker genes nor on lipid accumulation, both under basal conditions and under β-adrenergic stimulation. Likewise, there was no effect on lipid accumulation in the subsequent gain- and loss-of-function experiments in mature brown adipocytes. On transcriptome level, we hypothesised that gain- and loss-of function of *Gm15551* should lead to opposite effects on the gene expression of potential target genes of *Gm15551*. However, there were no genes that were significantly regulated in both datasets. Furthermore, most genes differentially regulated in one dataset did not even show a (non-significant) regulation in the other one. Those genes that were oppositely regulated by *Gm15551* gain- and loss-of-function such as *Lcn2*, *Saa3*, and *Hp* are inflammatory markers (Maffei et al., 2016; Sommer et al., 2009, 2008). We have previously found them to be differentially regulated by other sgRNAs in several datasets using the wt1-SAM model system and therefore interpret them as a model specific artefact.

As impaired iBAT function has been shown to render mice susceptible to diet-induced obesity and insulin intolerance (Guerra et al., 2001; Lowell et al., 1993), we put ∆Gm15551 mice on HFD for 12 weeks and additionally repeatedly tested their response to cold treatment by indirect calorimetry. While both the prolonged HFD treatment and sex caused clear differences, we could not detect any significant effect of the *Gm15551* loss-of-function on the examined adipose tissue and metabolic parameters such as energy expenditure, body weight and glucose tolerance. However, as the dataset is currently not well balanced, several of the experiments could so far not be statistically analysed. The last two cohorts are expected to be analysed in early 2022. When we analysed adipose tissue transcriptomes, clear differences between the white and brown adipose tissues as well as between iBAT from cold treated and control animals became evident. However, the *Gm15551* knockout only caused a minor number of differentially expressed genes, not exceeding what is expected as false positives.

We have identified a set of adipose tissue specific, HFD and cold regulated lncRNAs from which we characterised *Gm15551*. Albeit it is highly upregulated during brown adipogenesis and its expression correlates with iBAT activity, we could not detect a phenotype of either gain or loss-of-function of *Gm15551 in vitro*. Likewise we could not detect a detect a *Gm15551* related phenotype when performing comprehensive transcriptomic and adipose tissue physiologic phenotyping *in vivo*. In conclusion, our findings indicate that *Gm15551* is dispensable for iBAT development and function, despite its marked upregulation during initial adipose tissue development. This result is in concordance with a study, which previously found *Gm15551* to be upregulated in both white and brown adipogenesis, but detected no effect of small interfering RNA (siRNA) mediated knockdown of *Gm15551* on white adipocyte differentiation (Sun et al., 2013). While we have not ruled out a potential effect of a knockdown of *Gm15551* in brown preadipocytes on adipogenesis, the lack of a phenotype in white adipogenesis as well as *in vivo* investigations makes it appear implausible to find a phenotype in brown adipogenesis. Functional redundancy has been reported for coding genes such as CD34 and for duplicated genes in general (Hughes et al., 2020; Qian et al., 2010). Furthermore, lncRNAs have been shown to have tissue and cell state dependent and potentially very subtle functions (Derrien et al., 2012). We cannot rule out the possibility of the existance of other genes showing functional redundancy to *Gm15551* hiding any effects of the *Gm15551* loss-of-function. It was recently reported that in mouse and zebra fish several lncRNAs, which were selected because of their high expression levels, conservation or because they were located proximal to known coding developmental regulatory genes, had no effect on embryogenesis, viability and fertility (Goudarzi et al., 2019; Han et al., 2018). In conclusion, while *Gm15551* is specifically expressed in adipose tissues and we subjected mice both to HFD and cold treatment, two major stressors of adipose tissue (Alcalá et al., 2017; Sanchez-Gurmaches et al., 2016), it is possible that *Gm15551* exhibits either a very subtle function undetectable by our measurements, a context-dependent function in a specific cellular state that we have not investigated, or it might also be that *Gm15551* has no biological function in murine adipose tissue.

## 7 Material and methods

### Animal experiments

Unless otherwise stated, mice were kept at 22 °C to 24 °C on a regular 12 h light cycle with *ad libitum* access to food and water. Wild type C57BL/6N mice used for the detection of differentially regulated genes and ChIP-Seq in iBAT were fed chow diet (Ssniff V1554) up to the age of 8 weeks, where the respective cohorts were put on HFD (Ssniff D12492 (I) mod.) for 12 weeks and kept at 4 °C for 24 h. ΔGm15551 mice used for transcriptomics analyses were fed chow diet (Altromin 1324). ΔGm15551 mice used for metabolic phenotyping were fed chow diet (Altromin 1314). Respective cohorts were put on high fat diet (Ssniff D12492 (I) mod.) for 12 weeks starting from 8 weeks of age.

### Generation of *Gm15551* knock out animals

The mouse model for genetic deficiency of *Gm15551* was generated at the Czech Centre for Phenogenomics using CRISPR/Cas9 targeting exon 1 of *Gm15551* on the background of C57BL/6N. Sanger sequencing verified a 1113 bp deletion including exon 1 of *Gm15551* (3:126462197-126463309, GRCm38). Knock out mice were backcrossed with wild type animals for two generations to minimise the risk of off target effects. Genotyping was carried out by two separate PCRs using primers F3/R2 (Tab S1; 372 bp and 1377 bp for knock out and wild type alleles respectively) and F5/R4 (234 bp for wildtype allele only). DNA extracted from tail tips using proteinase K digestion and Chelex 100 Resin (Biorad 1432832) was amplified for 30 cycles at 95 °C, 60 °C and 72 °C for 30 s each and visualised using capillary gel electrophoresis (Fragment Analyzer, Advanced Analytics).

### Indirect calorimetry

Prior to the experiment, a complete calibration protocol for the gas analysers was run according to the manufacturer’s recommendations and mice were weighed. The mice were singly housed in a PhenoMaster device (TSE systems) at a regular 12 h light cycle and 55 % relative humidity with *ad libitum* access to water and the respective diet. At 11, 15 and 19 weeks of age, mice underwent a temperature challenge starting at 23 °C, followed by 6 h at 30 °C, 18 h at 4 °C and 9 h at 23 °C again. Sampling rate was 15 min.

### IPGTT

Animals were fasted overnight (16 h to 18 h) with free access to water. After weighing, the mice received 2 g kg^−1^ i.p. glucose. Blood glucose was measured fasted and after 15, 30, 60 and 120 min using a standard glucometer.

### Adipocyte diameter

Hematoxylin and eosin staining was performed on tissue slices and slides were scanned. Two representative areas per tissue were exported and analysed using adiposoft (Galarraga et al., 2012).

### RNA isolation and reverse transcription

Cells or frozen tissue samples were homogenised and lysed in TRIsure (Bioline). Total RNA was isolated using EconoSpin All-In-One Mini Spin Columns (EconoSpin 1920-250) and reverse transcribed into cDNA using the High Capacity cDNA Reverse Transcription Kit (Applied Biosystems 4368814) following the manufacturer’s instructions.

### Quantitative polymerase chain reaction (qPCR)

qPCR was performed in 384 well format in a LightCycler 480 II (Roche). 4 μl of 1:20 diluted cDNA, 0.5 μl gene specific primer mix (5 μM each) and 4.5 μl FastStart Essential cDNA Green Master (Roche) were amplified using 45 cycles of 25 s at 95 °C, 20 s at 58 °C and 20 s at 72 °C after 300 s at 95 °C initial denaturation. All combinations of primers and samples were run in duplicates and *C*_*q*_ values calculated as the second derivative maximum. Genes of interest were normalised against housekeeper genes using the Δ*C*_*q*_ method. The primers used in this study can be found in Tab S4.

### Total RNA sequencing

RNA sequencing and library preparation were performed at the Cologne Center for Genomics (Cologne, Germany) according to their standard protocols. Before RNA sequencing, rRNA was depleted according to the instructions of the Illumina TruSeq kit. All sequencing experiments were accomplished with a paired-end protocol and a depth resulting in 50 *×* 10^6^ to 75 *×* 10^6^ paired reads per sample. Before RNA sequencing, genomic DNA was eliminated following the instructions of the TURBO DNA-freeTM Kit and subsequently 1 μl RNA was used to examine RNA integrity in an Agilent 2100 Bioanalyzer Analysis System.

### Poly A RNA sequencing

Paired end libraries were constructed using the NEBNext Ultra II RNA Library Prep Kit for Illumina following the manufacturer’s protocol and sequenced on an Illumina NovaSeq 6000 in 2 x 50-bp paired end reads.

### RNA sequencing data analysis

Reads were quality filtered using cutadapt (Martin, 2011). For visualisation, reads were mapped to the GRCm38 genome using STAR (Dobin et al., 2013). For quantification, reads were mapped to the Gencode M22 transcriptome or a combination of M22 and RNAcentral 5 using salmon (Patro et al., 2017).

### ChIP sequencing

For histone modification sequencing, brown adipose tissues of two mice were used each sequencing experiment. Prior to ChIP, BAT was dissociated using a gentleMACSTM Dissociator (Miltenyi biotec, Germany). The cell suspension was cross linked with 1 % formaldehyde for 10 min at RT and the reaction was quenched with 0.125 M glycine for 5 min to 10 min at RT. Cells were washed twice with cold PBS and PMSF and snap-frozen in liquid nitrogen before storing at −80 °C.

Frozen pellets were thawed on ice for 30 min to 60 min. Pellets were resuspended in 5 ml lysis buffer 1 (50 mM Hepes, 140 mM NaCl, 1 mM EDTA, 10 % glycerol, 0.5 % NP-40, 0.25 % Triton X-100) by pipetting and then rotated vertically at 4 °C for 10 min. Pellets were resuspended in 5 ml lysis buffer 2 (10 mM Tris, 200 mM NaCl, 1 mM EDTA, 0.5 mM EGTA) and incubated at vertical rotation and at room temperature for 10 min. Samples were centrifuged for 5 min at 1350 g at 4 °C and supernatant was carefully aspirated. Then, samples were resuspended in 3 ml lysis buffer 3 (10 mM Tris, 100 mM NaCl, 1 mM EDTA, 0.5 mM EGTA, 0.1 % Na-deoxycholate, 0.5 % N-lauroylsarcosine) and were separated into 2 times 1.5 ml in 15 ml polypropylene tubes, in which they were sonicated with the following settings by Bioruptorő Plus sonication: power = high, on interval = 30 s, off interval = 45 s, total time = 10 min (18 cycles of on/off). Sonicated samples were transferred to a 1.5 ml microfuge tube and were centrifuged for 10 min at 16 000 g at 4 °C to pellet cellular debris. 10 % of sample solution were stored to be used as input control, while the rest was used for ChIP.

To capture different histone modifications, 5 μg to 10 μg of the respective antibodies (Tab S5) were added to the sonicated ChIP reaction and rotated vertically at 4 °C overnight. 100 μl Dynabeads (Protein A or Protein G) for each ChIP sample were prepared according to the manufacturer’s instructions, mixed with 1 ml of antibody-bound chromatin and rotated vertically at 4 °C for at least 2 h to 4 h. Bound beads were washed at least five times in 1 ml cold RIPA (50 mM Hepes, 500 mM LiCl, 1 mM EDTA, 1 % NP-40, 0.7 % Na-deoxycholate) and once in 1 ml cold TE buffer containing 50 mM NaCl. Samples were eluted for 15 min with elution buffer (50 mM Tris, 10 mM EDTA, 1 % SDS) at 65 °C and continuously shaken at 700 min^−1^. Beads were separated using a magnet and 200 μl supernatant were transferred to fresh microfuge tubes. Input samples were thawed and mixed with 300 μl elution buffer. ChIP/input samples were incubated at 65 °C in a water bath overnight to reverse the cross linking reaction. TE buffer was added at room temperature to dilute SDS in both ChIP and input samples. For digestion of RNA and protein contamination, RNase A was added to the samples and incubated in a 37 °C water bath for 2 h; then proteinase K was added to a final concentration of 0.2 mg ml^−1^ and incubated in a 55 °C water bath for 2 h. Finally, DNA was extracted using a standard phenol-chloroform extraction method at room temperature and DNA concentrations were measured using a NanoDrop ND-1000 spectrophotometer or Qubit dsDNA HS Assay Kit and stored at −80 °C until sequencing.

### ChIP sequencing data analysis

Reads were mapped to the GRCm38 genome using bowtie2 (Langmead and Salzberg, 2012) after quality filtering by cutadapt.

### Tissue specificity

Tissue specificity scores were calculated for every gene over seven metabolically active tissues as 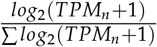 as described by (Alvarez-Dominguez et al., 2015) from RNA-Seq data that we have previously published (GEO: GSE121345.

### Gene set enrichment analysis

Gene set enrichment was done using topGO for GO (Alexa et al., 2006) and ReactomePA for reactome (Yu and He, 2016).

### Assessment of coding potential

Coding potential was calculated using CPAT (Wang et al., 2013). Ribosome scores were calculated as log2(TRAP/totalRNA), where TRAP are the normalised counts from a publicly available dataset of translating ribosomal affinity purification (TRAP) of mouse iBAT (GEO: GSE103617) and totalRNA are the normalised counts from the room temperature control diet iBAT total RNA samples.

### Primary cell culture

Inguinal and epididymal white as well as intrascapular brown adipose tissues from 6 to 8 week old C57BL/6J mice were dissected, minced and digested with collagenase II (worthington) and dispase II (Sigma, iBAT only). Cells were seeded in 24 well plates and grown in DMEM/Ham’s F12 medium supplemented with 0.1 % Biotin/D-Pantothenate (33 mM/17 mM), 1 % penicillin-streptomycin and 20 % FBS. Upon reaching confluency, FBS concentration was reduced to 10 % and differentiation was induced using 1 μM rosiglitazone, 850 nM insulin, 1 μM dexamethasone, 250 μM 3-isobutyl-1-methylxanthin (IBMX), 125 μM indomethacine (brown only) and 1 nM triiodothyronine (T3) (iBAT only). Subsequently, medium was changed every other day for medium containing 10 % FBS, rosiglitazone and T3 (iBAT only). Full differentiation was reached 7 days after induction. Cells were stimulated using 1 μM isoproterenol or 10 μM CL316243.

### Cultivation of brown adipocyte cell lines

The wt1-SAM brown preadipocyte cell line was a kind gift from Dr. Brice Emanuelli. The cells were grown in high glucose DMEM supplemented with 10 % FBS and 1 % penicillin-streptomycin. After reaching confluency, differentiation was induced by 0.5 μM rosiglitazone, 1 nM T3, 1 μM Dexamethasone, 850 nM insulin, 125 μM indomethacine and 500 μM IBMX. Two days later, medium was exchanged for medium supplemented with 0.5 μM rosiglitazone and 850 nM insulin. Afterwards, medium was changed for medium containing 0.5 μM rosiglitazone every second day. Full differentiation was reached 7 days after induction.

PIBA cells were cultured in the same medium as wt1-SAM cells. A common induction/stimulation cocktail consisting of 10 μM rosiglitazone, 1 nM T3, 0.5 μM Dexamethasone, 850 nM insulin, 12.5 μM indomethacine and 125 μM IBMX was used to differentiate cells.

### *In vitro* gain and loss of function studies

For *in vitro* gain of function studies using the wt1-SAM cell line, sgRNAs were designed using CRIS-Pick (Doench et al., 2014) and cloned into the sgRNA(MS2) cloning backbone (addgene 61424) as described by Konermann et al. (2015). Empty vector and *Ucp1* targeting sgRNAs were used as controls and were kind gifts of Dr. Brice Emanuelli (Lundh et al., 2017). Target sequences used are found in Tab S2.

LNA gapmers designed and synthesized by Qiagen were used for *in vitro* loss of function experiments. Two non targeting scrambled LNAs were used as control (Tab S3).

Preadipocytes were transfected by seeding 30 000 cells per well of a 24 well plate in growth medium and adding 1.5 μl TransIT-X2 (Mirus) and 125 ng plasmid DNA or 1.4 μl LNA (10 μM) in 50 μl OptiMEM I once the cells had attached.

In order to transfect mature adipocytes, 3 μl TransIT and 250 ng plasmid DNA or 1.4 μl (10 μM) in 100 μl Opti-MEM I were pipetted into a well of a 24 well plate. After 15 min, 500 000 cells resuspended in 500 μl Opti-MEM were added. 24 h later, medium was changed for regular differentiation medium.

### Oil red O staining

Cells were fixed with 4 % formalin for 30 min, rinsed once with water followed by 60 % isopropanol. Cells were stained with Oil Red O (0.3 % in 60 % isopropanol) for 10 min. Excess dye was rinsed with water. For quantification, the Oil Red O was eluted in 100 % isopropanol and OD measured at 520 nm in a multi plate reader.

### Statistical analysis

Statistics for RNA-Seq data was done in DESeq2 (Love et al., 2014) using LRTs for factors with multiple levels or for analysing multiple factors at once and wald tests otherwise. Log fold changes were shrunken and *s*-values calculated using apeglm (Zhu et al., 2018). Data from animal experiments with repeated measurements were averaged over temperature and day/night conditions and analysed using mixed-effects models with the body weight as cofactor and individual animal as random variable. Other data was analysed using Student’s *t*-tests and adjusted for multiple testing using Holm’s method. Shown are individual values in addition to mean *±*standard error of the mean.

## 3 Abbreviations

ANCOVA: analysis of covariances
ANOVA: analysis of variances
ATP: adenosine triphosphate
BAT: brown adipose tissue
cDNA: complementary DNA
ChIP: chromatin immunoprecipitation
DMEM: Dulbecco’s modified Eagle’s medium
DNA: deoxyribonucleic acid
eRNA: enhancer RNA
eWAT: epididymal white adipose tissue
FBS: fetal bovin serum
GO: gene ontology
HFD: high fat diet
iBAT: interscapular brown adipose tissue
IBMX: 3-isobutyl-1-methylxanthin
IPGTT: intraperitoneal glucose tolerance test
iWAT: inguinal white adipose tissue
LNA: locked nucleic acid
log2FC: log2 fold change
lncRNA: long non-coding RNA
LRT: likelihood ratio test
miRNA: micro RNA
qPCR: quantitative polymerase chain reaction
RNA: ribonucleic acid
SDS: Sodium dodecyl sulfate
sgRNA: single guide RNA
siRNA: small interfering RNA
T3: triiodothyronin
TRAP: translating ribosomal affinity purification
WAT: white adipose tissue

## 8 Acknowledgements

CH was funded by the National Taiwan University and the grant 109-2917-I-002-029 from the Ministry of Science and Technology, Taiwan. The authors used services of the Czech Centre for Phenogenomics supported by the Czech Academy of Sciences RVO 68378050 and by the project LM2018126 Czech Centre for Phenogenomics provided by Ministry of Education, Youth and Sports of the Czech Republic. JWK and CHE were funded by the University of Southern Denmark and the Danish Diabetes Academy, which is in turn funded by the Novo Nordisk Foundation.

## 9 Author contributions

CHE performed experiments with primary adipocytes, *Gm15551* gain of function experiments in preadipocytes, bioinformatic analyses, analysed the data, designed the study and wrote the manuscript. CH performed the *Gm15551* gain and loss of function experiments in mature adipocytes and helped with the ΔGm15551 loss of function experiments for transcriptomics. SK performed the experiments with wild type mice used for transcriptomics and performed ChiP. PK generated the ΔGm15551 knockout mouse model. JP, JR and DPR planned and performed the metabolic phenotyping of the ΔGm15551 mice. RS supervised the generation and metabolic phenotyping of the ΔGm15551 mice. JWK designed and supervised the study and wrote the manuscript.

## 14 Supplementary tables

**Table S1:**
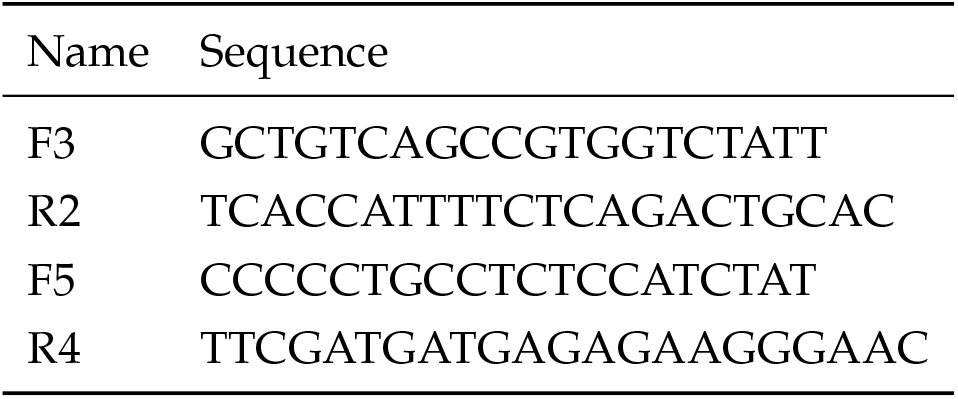
Sequences of genotyping PCR primers.

**Table S2:**
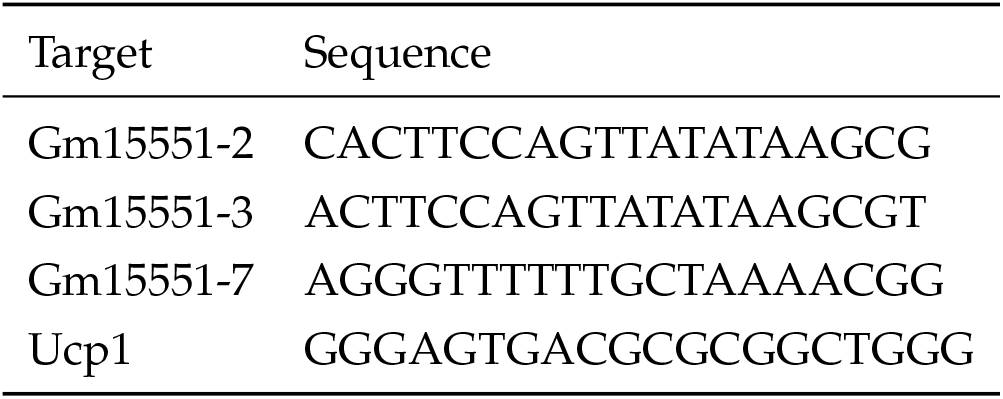
Sequences of sgRNAs.

**Table S3:**
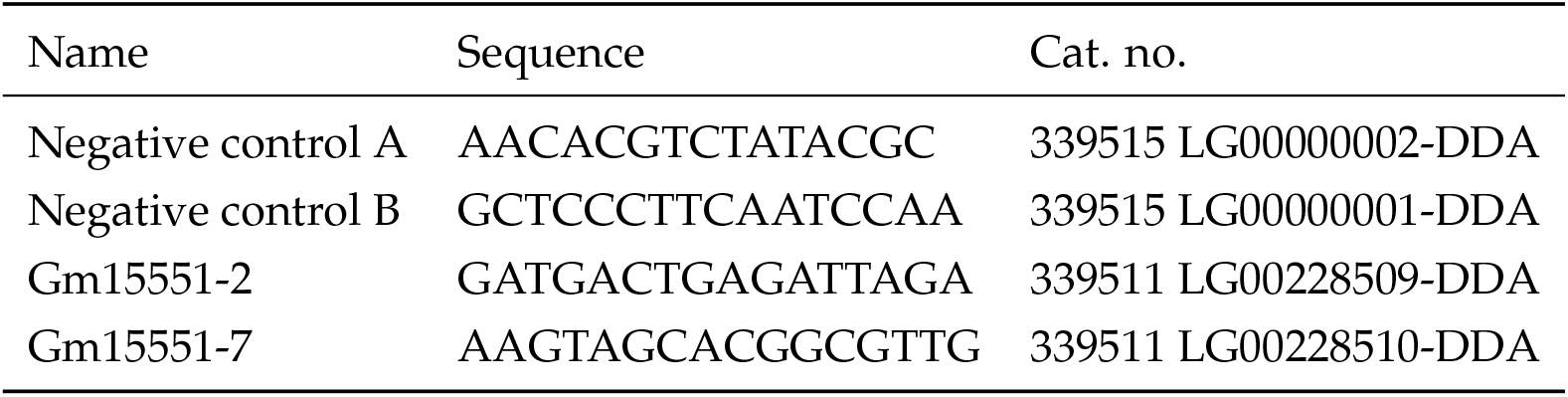
Sequences of LNAs.

**Table S4:**
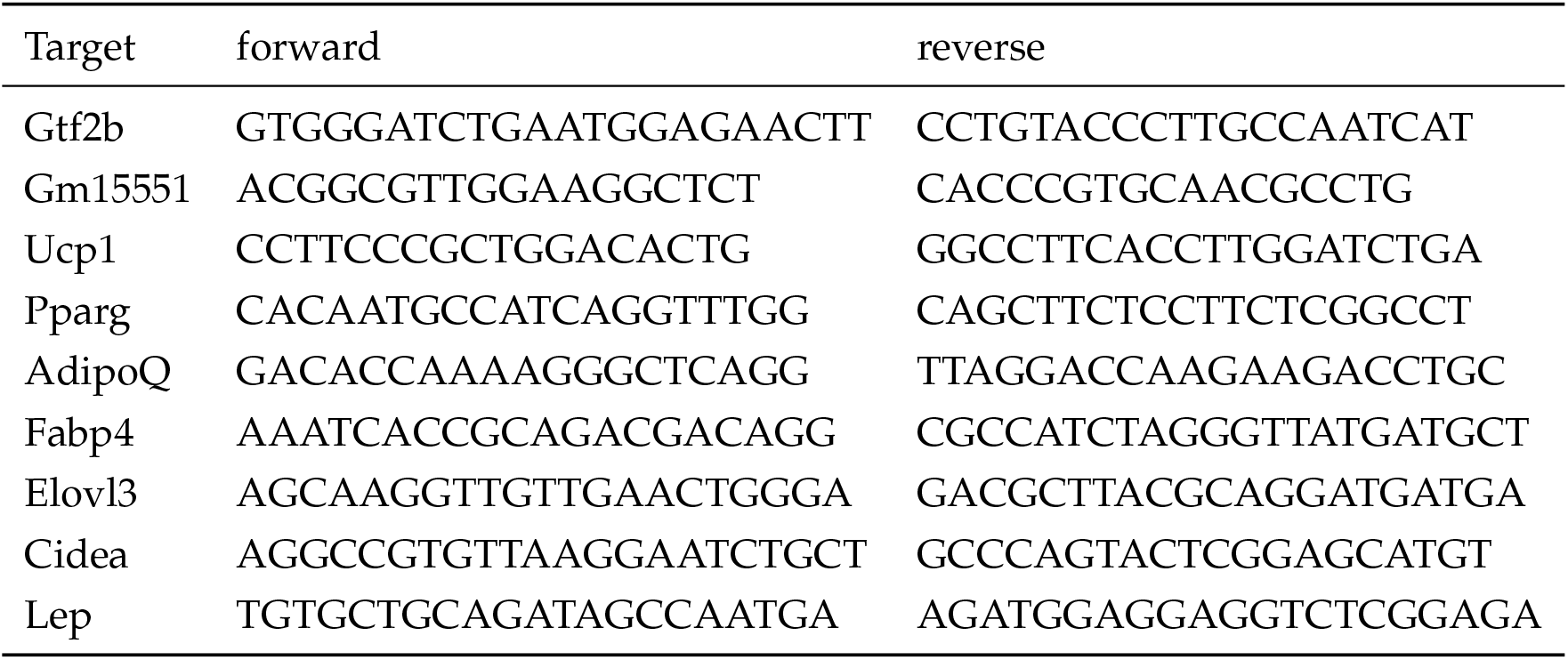
Sequences of qPCR primers.

**Table S5:** Antibodies used for chromatin modification ChIP-Seq.

| Target | Source | Dilution |
| --- | --- | --- |
| Histone H3K27ac | Active Motif 39133 | 5 µl |
| Histone H3K27me3 | Active Motif 39155 | 5 µl |
| Histone H3K4me1 | Active Motif 39297 | 10 µl |
| Histone H3K4me3 | Active Motif 39159 | 3 µl |

## 15 Legends

**Fig S1:**
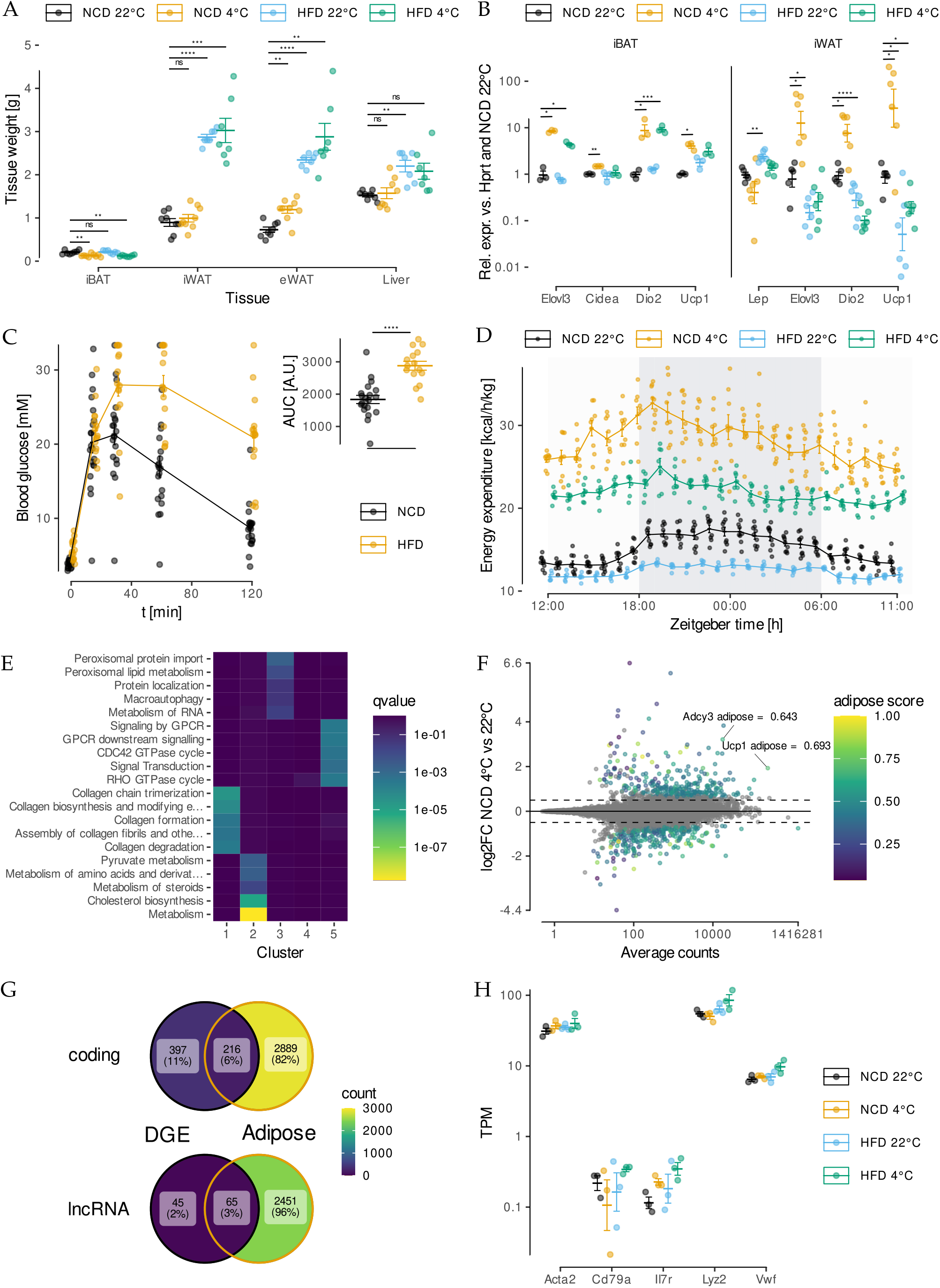
RNA-Seq reveals temperature and obesity dependent changes in iBAT lncRNA expression. **A** Adipose tissues and liver weights of 20 week old mice after cold and/or HFD treatment (*t*-test, n = 7 - 9). **B** Expression of common and brown specific adipose marker genes and the macrophage marker *Emr1* in iBAT and iWAT of cold and/or HFD challenged mice (*t*-test, n = 3 - 6). **C** IPGTT of HFD and control diet fed animals at 12 weeks to 14 weeks of age (*t*-test, n = 15). **D** Energy expenditure of HFD or control animals kept at either 22 °C or 4 °C measured by indirect calorimetry (n = 5 - 8). **E** Reactome pathway enrichment analysis for the clusters in Fig1 A. **F** Expression levels and changes for coding genes in iBAT from cold treated compared to control mice. Genes showing significant differential gene expression are colour coded indicating their adipose tissue specificity (wald test, n = 6, *s* < 0.05, *H*_0_: log2FC > 0.5). **G** Overlap of differential gene expression (wald test, *s* < 0.05) and adipose tissue specificity (adipose score > 0.5) for coding and lncRNA genes. **H** Expression levels of immune cell marker genes in the total transcriptomes from iBAT of cold and/or HFD treated mice.

**Fig S2:**
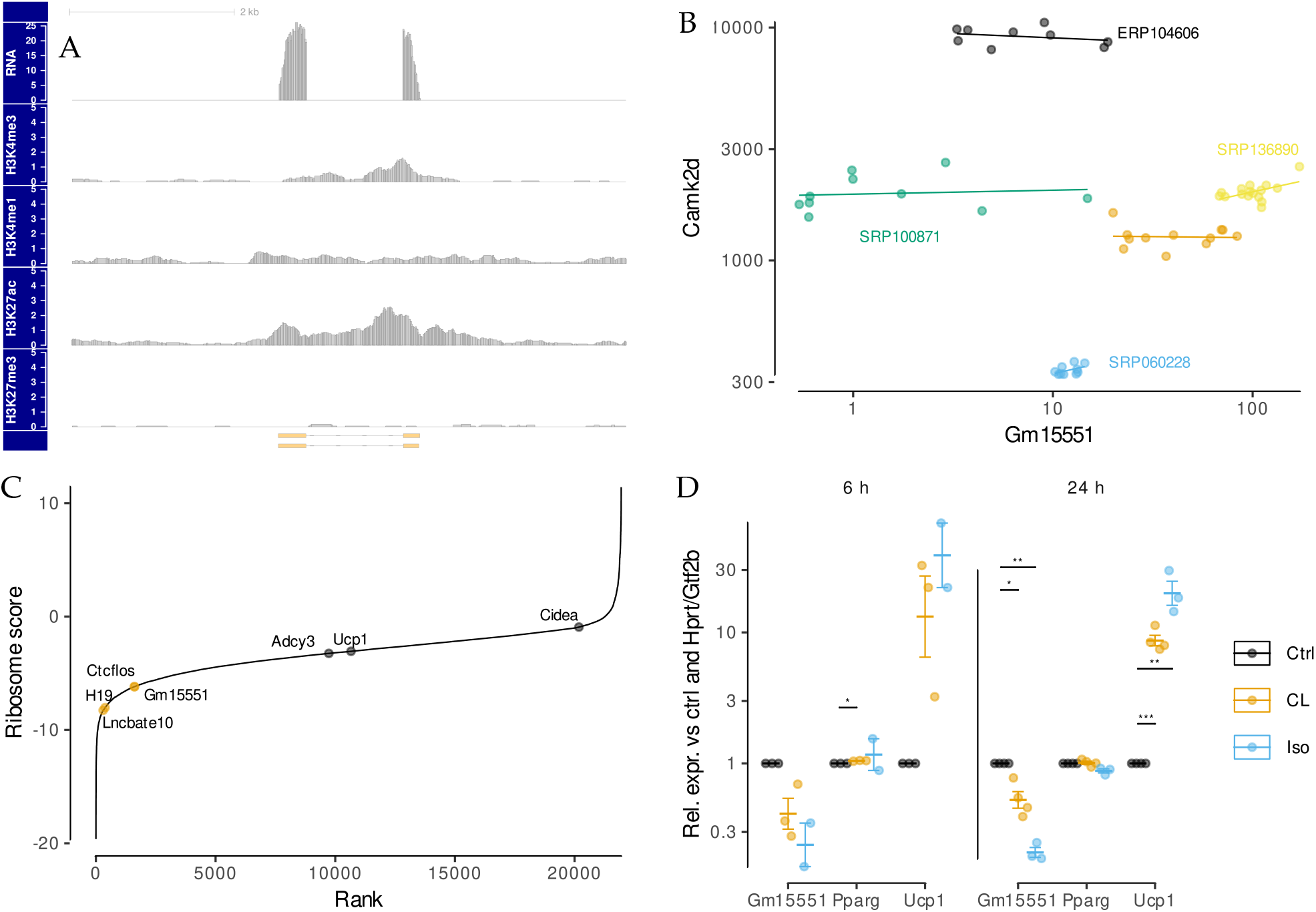
*Gm15551* is an adipose tissue specific, diet and temperature regulated lncRNA. **A** Genomic locus of *Gm15551* showing the RNA expression as well as chromatin modifications in iBAT of control mice. **B** Expression of *Gm15551* vs. *Camk2d* in indicated public RNA-Seq data sets. (Orange is the dataset from this study.) **C** Ranked ribosome scores as calculated from TRAP-Seq of murine iBAT (PRJNA402074). **D** Expression of *Gm15551*, *Pparg* and *Ucp1* in fully differentiated PIBA cells, stimulated with isoproterenol or CL316243 for either 6 h or 24 h (paired *t*-test, n = 2-3).

**Fig S3:**
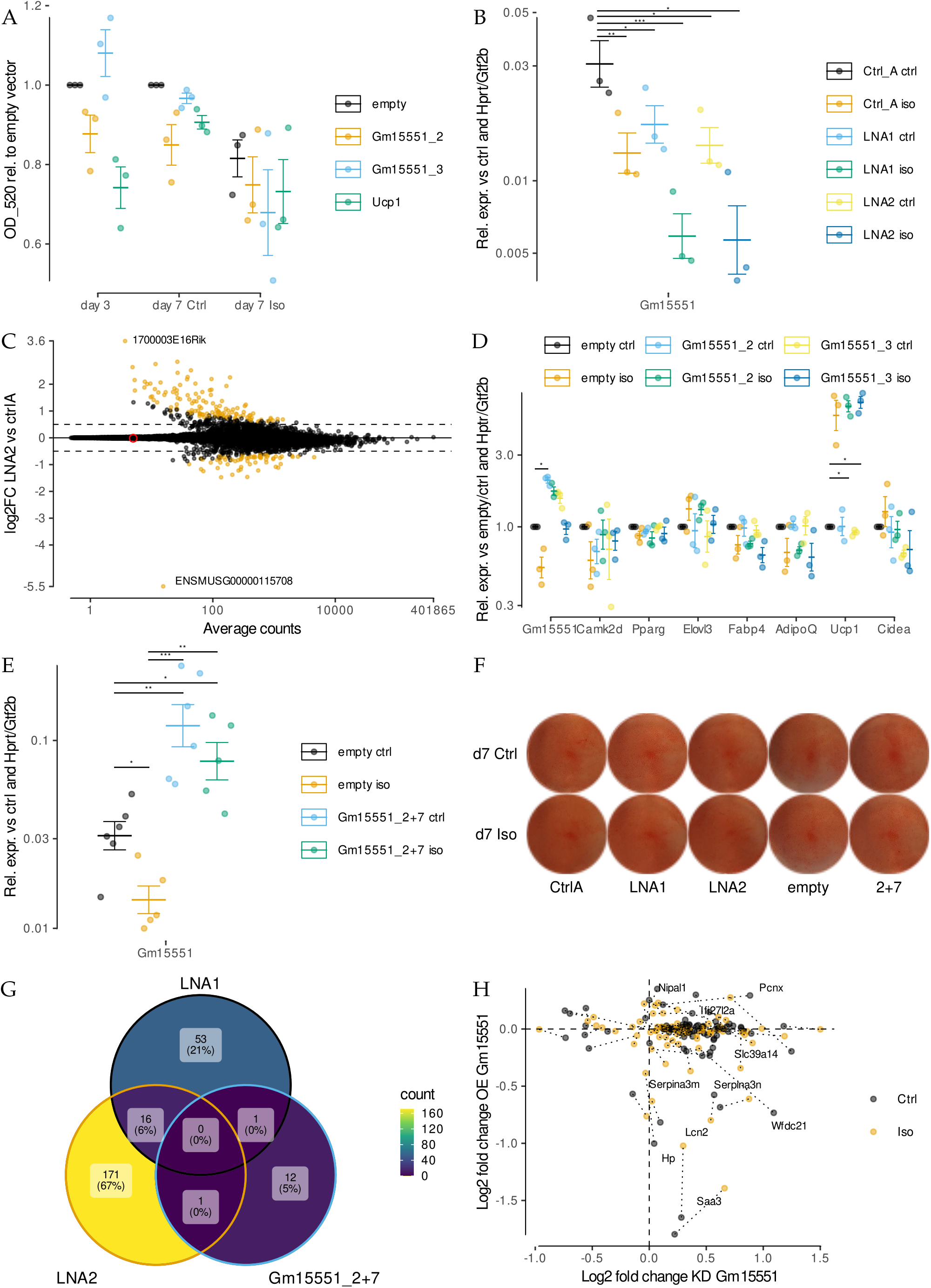
*Gm15551* is dispensable for iBAT function *in vitro*. **A** Quantification of lipid accumulation by oil red O staining of wt1-SAM cells transfected with plasmids encoding sgRNAs targeting *Gm15551* or empty vector two days before induction of differentiation (paired *t*-test, n = 3 - 6). **B** Efficiency of the knockdown of *Gm15551* using wo different LNAs in mature wt1-SAM cells (paired *t*-test, n = 3). **C** Effect of knockdown of *Gm15551* using LNA2 at day 4 of differentiation on gene expression in mature wt1-SAM cells (wald test, log2FC > 0.5, n = 6, *s* < 0.05). **D** Gene expression of *Gm15551*, *Camk2d*, the general adipocyte markers *Pparg*, *Elovl3*, *Adipoq* and *Fabp4* as well as the brown adipocyte marker genes *Ucp1* and *Cidea* in mature wt1-SAM cells after overexpression of *Gm15551* at day 4 of differentiation (paired *t*-test, n = 3). **E** Efficiency of overexpression of *Gm15551* in mature wt1-SAM cells using a combination of 2 sgRNAs (wald test, log2FC > 0.5, n = 6, *s* < 0.05). **F** Oil red O staining of mature wt1-SAM cells after transfection with LNAs or plasmids encoding sgRNAs targeting *Gm15551* at day 4 of differentiation. **F** Effect of gain and loss of function of *Gm15551* in mature adipocytes on lipid accumulation. **G, H** Overlap between genes showing differential regulation and comparison of their gene expression changes in wt1-SAM cells upon knockdown or overexpression of *Gm15551* (*s* < 0.05) at day 4 of differentiation.

**Fig S4:**
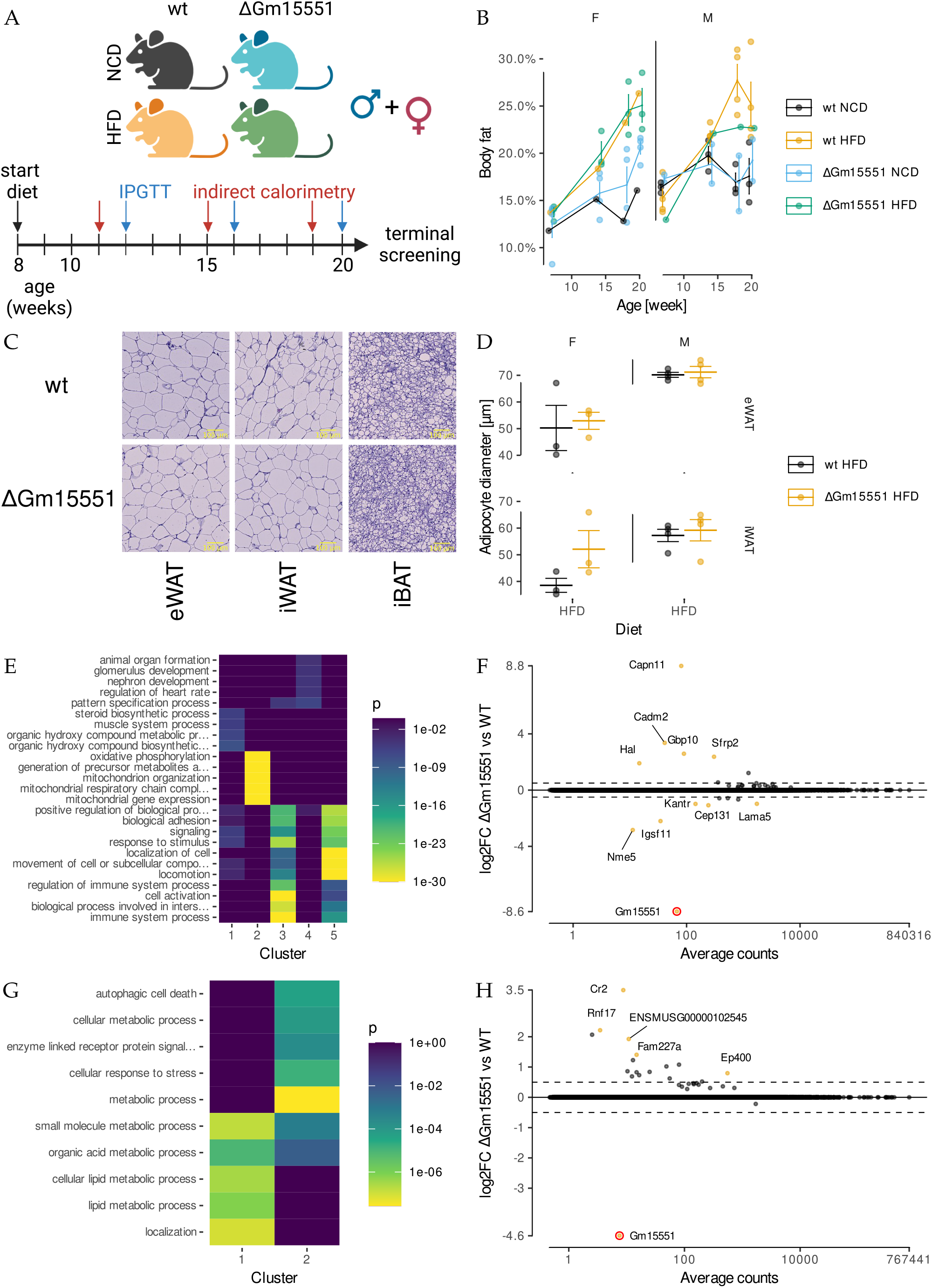
*Gm15551* is dispensable for iBAT function *in vivo*. **A** Experimental design. Male and female ΔGm15551 and wild type litter mates were either put on HFD or control diet for 12 weeks starting at 8 weeks of age. IPGTT and indirect calorimetry measurements were repeatedly performed at the indicated timepoints. Created with BioRender.com. **B** Body fat percentage of ΔGm15551 and wild type mice fed a high fat or control diet. **C, D** Representative microphotographs (C) and adipocyte diameters (D) in different adipose tissue from 20 week old wt and ΔGm15551 mice fed a HFD. **E** GO enrichment analysis for the gene clusters shown in Fig 4E. **F** Gene expression changes induced by the knockout of *Gm15551* in different adipose tissues (wald test, n = 3, *s* < 0.05, *H*_0_: log2FC > 0.5). **G** GO enrichment analysis for the gene clusters shown in Fig 4F. **H** Gene expression changes induced by the knockout of *Gm15551* in iBAT of cold treated and room temperature housed mice (wald test, n = 3 or 5, *s* < 0.05, *H*_0_: log2FC > 0.5).

